# Nocturnal pollination services as an ecological safety net for papaya production in a tropical biodiversity hotspot

**DOI:** 10.64898/2026.07.17.739175

**Authors:** Mark Otieno, Marcell K. Peters, Sara Marie Plesser, Jie Zhang, Ingolf Steffan-Dewenter

**Affiliations:** Department of Animal Ecology and Tropical Biology, Biocenter, University of Würzburg, Am Hubland, 97074, Würzburg, Germany; Animal Ecology Group, BIOM, University of Bremen, James-Watt-Straße 1, 28359 Bremen, Germany; Department of Water and Agricultural Resource Management, University of Embu, P.O. Box 6-60100, Embu, Kenya

## Abstract

Pollination is of key importance for tropical agriculture, yet a bee-centred perspective often overlooks the critical role of nocturnal insects in pollination services. We evaluated the relative contributions of diurnal and nocturnal pollinators to papaya yields on subsistence farms in the Taita Hills, Kenya, a globally important biodiversity hotspot. We combined flower visitor observations with five experimental treatments: open pollination, wind-only (closed), hand pollination, and temporal exclusions for day-only and night-only services. Our results reveal complete temporal niche partitioning between diurnal bees and nocturnal hawkmoths (Sphingidae). We found that diurnal and nocturnal pollinators provide additive, rather than redundant, services, contributing to papaya fruit set and weight to a similar extent. Despite these contributions, supplementary hand-pollination experiments indicated a significant yield gap (36.9% lower fruit set), suggesting substantial pollinator limitation. While landscape-scale factors had no significant effects, nocturnal pollinators responded positively to organic management, whereas diurnal visitors did not. This indicates that local habitat quality and ecological intensification are more critical for sustaining specialised pollinators than landscape composition. Our study highlights the vital role of nocturnal insects as an ecological safety net and emphasises that their conservation through local organic practices is essential for securing yield quantity and quality in tropical papaya farming systems.

## Introduction

Tropical agroecosystems are facing unprecedented pressure to scale up production, driven by a surging global population and expanding international trade networks [1]. Because many tropical regions feature highly weathered soils and are highly susceptible to biodiversity loss, agricultural productivity in these biomes remains closely linked to ecosystem services, such as nutrient cycling, natural pest control, and biotic pollination provided by wild species [2,3]. As conventional intensification increasingly threatens the landscape-level biodiversity that underpins these ecological functions [4], transition frameworks such as agroecology and organic management have become central to achieving sustainable crop production [1]. Within this broader tropical agricultural landscape, high-value horticultural commodities are experiencing a pronounced shift. Notably, the global demand for tropical fruits, specifically papaya (*Carica papaya* L.), has risen rapidly in recent decades, driven by growing consumer awareness of its nutritional value and the expansion of trade networks [5,6].

Across Sub-Saharan Africa, smallholder fruit production plays a significant role in rural livelihoods by generating household income and providing essential micronutrients, such as vitamins A and C, directly from on-farm production [7,8]. Because these small-scale horticultural systems often operate under low-input regimes, their productivity remains closely tied to the ecosystem services regulating wild, diverse pollinator communities [9,10]. However, landscape degradation and conventional intensification increasingly threaten the wild insect populations that underpin these yields [11].

While the role of diurnal insects, particularly bees, in sustaining agricultural yields is well documented, the vital contributions of nocturnal pollinators, such as moths and bats, remain critically understudied in tropical agroecosystems [13,14]. This knowledge gap is particularly problematic because nocturnal plant-pollinator networks are often highly specialised and extremely sensitive to anthropogenic disturbances, such as chemical pesticide use, habitat loss and fragmentation [15]. Ecological intensification, through strategies such as organic farming, offers a viable pathway to mitigate these pressures by enhancing local floral resources and eliminating synthetic inputs, thereby potentially stabilising both daytime and nighttime ecosystem services [16,17]. However, despite the clear theoretical benefits of these sustainable management frameworks, empirical assessments rarely examine how diurnal and nocturnal pollinator communities respond to organic management in tropical landscapes [14].

In *Carica papaya*, the path from flowering to a marketable harvest is shaped by its complex reproductive biology. This species is predominantly dioecious with individual plants bearing either male or female flowers [18]. This reproductive strategy makes the crop highly dependent on external biotic agents, principally insects, for pollen transfer among plants [19]. Without efficient inter-tree movement, the potential for high-yield harvests is diminished, making a study of pollination services relevant not merely as an academic pursuit but as a fundamental requirement for food security and farmers’ income [20].

This study assesses the individual and synergistic contributions of diurnal and nocturnal pollinators to fruit set and yields of *C. papaya* within the tropical agricultural mosaics of the Taita Hills in Kenya. To investigate the identified knowledge gaps, we addressed the following specific objectives:

1. Quantify daily activity patterns within the floral visitor community to determine the extent of temporal segregation between diurnal and nocturnal pollinators. This objective aims to uncover the baseline temporal dynamics of resource use within the pollinator community, providing insight into how temporal niche complementarity may stabilise plant-pollinator networks and mitigate competitive displacement.
2. Disentangle the contributions of diurnal and nocturnal pollinators to fruit set, fruit weight, and cumulative production through experimental pollination exclusion treatments. This objective allows us to quantify the yield gap associated with the pollinator limitation in specific temporal niches.
3. Evaluate how local management and landscape composition and configuration influence the abundance and community structure of diurnal versus nocturnal flower visitors. This objective aims to determine whether specialized nocturnal pollinators are more sensitive to local and landscape-scale management compared with diurnal pollinators, and to identify the most relevant spatial scales for sustaining pollination services.

## Results

### Pollen Limitation of Fruit Set and Weight

Our experimental manipulations revealed significant pollen limitations in both fruit quantity (number of fruits) and quality (fruit weight) across the study sites. Supplemental hand-pollination consistently outperformed natural open-pollination. For fruit set, supplemental pollination yielded an average of 6.62 fruits per tree, whereas open pollination yielded only 3.69 fruits per tree, representing a 36.9% gap in fruit set. A similar trend was observed for fruit weight. The average weight of fruits produced under supplemental pollination reached 1,329.43 g (±155.27 SE) per fruit, compared to 898.78 g (±132.30 SE) under open-pollination conditions. This translates to an average reduction of 430.65 g per fruit, resulting in a 32.4% reduction in total fruit weight directly attributable to pollen limitation.

### Papaya flower visitation

We recorded 2,056 floral visitors from seven insect orders and one avian group on papaya flowers at the 17 study sites. To characterise the floral visitor community, we quantified the mean abundance of various taxonomic and functional groups (Fig. 1). The pollinator community was heavily dominated by two primary groups: bees and hawkmoths, collectively accounting for approximately 89.3% of all observed flower visits. The remaining taxonomic groups, including beetles, flies, butterflies, ants, and birds, contributed marginally to total visitation rates, with mean abundances collectively falling below 10 individuals per site (Fig. 1).

**Figure 1:**
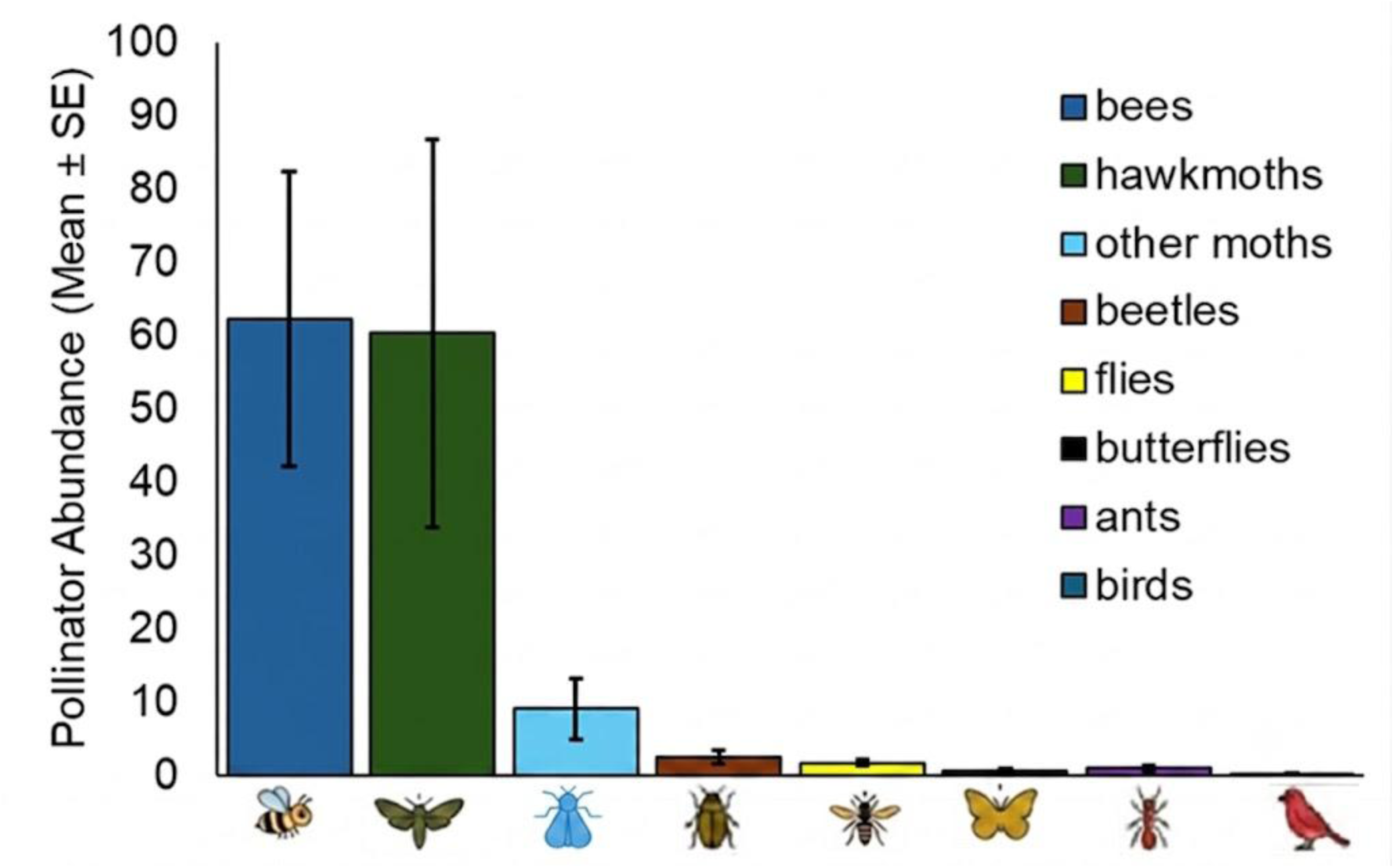
Mean abundance of eight dominant floral visitor taxa observed per papaya (*Carica papaya*) tree across distinct study sites and sampling periods. The percentages above the bars indicate the proportion of visits that occurred at night for each group. Data are derived from observations across 61 individual trees.

To characterise the pollination networks over time, we analysed visitation rates across different times of day and night (Fig. 2), revealing nearly complete temporal niche partitioning between bees and hawkmoths. Bee visitation was strictly diurnal, whereas hawkmoths served as the primary nocturnal vectors, exhibiting high nighttime activity and virtually no daylight presence (Fig. 2). Total floral visitation was significantly higher at night than during the day (z = –4.029, p < 0.001; Table 1).

**Figure 2:**
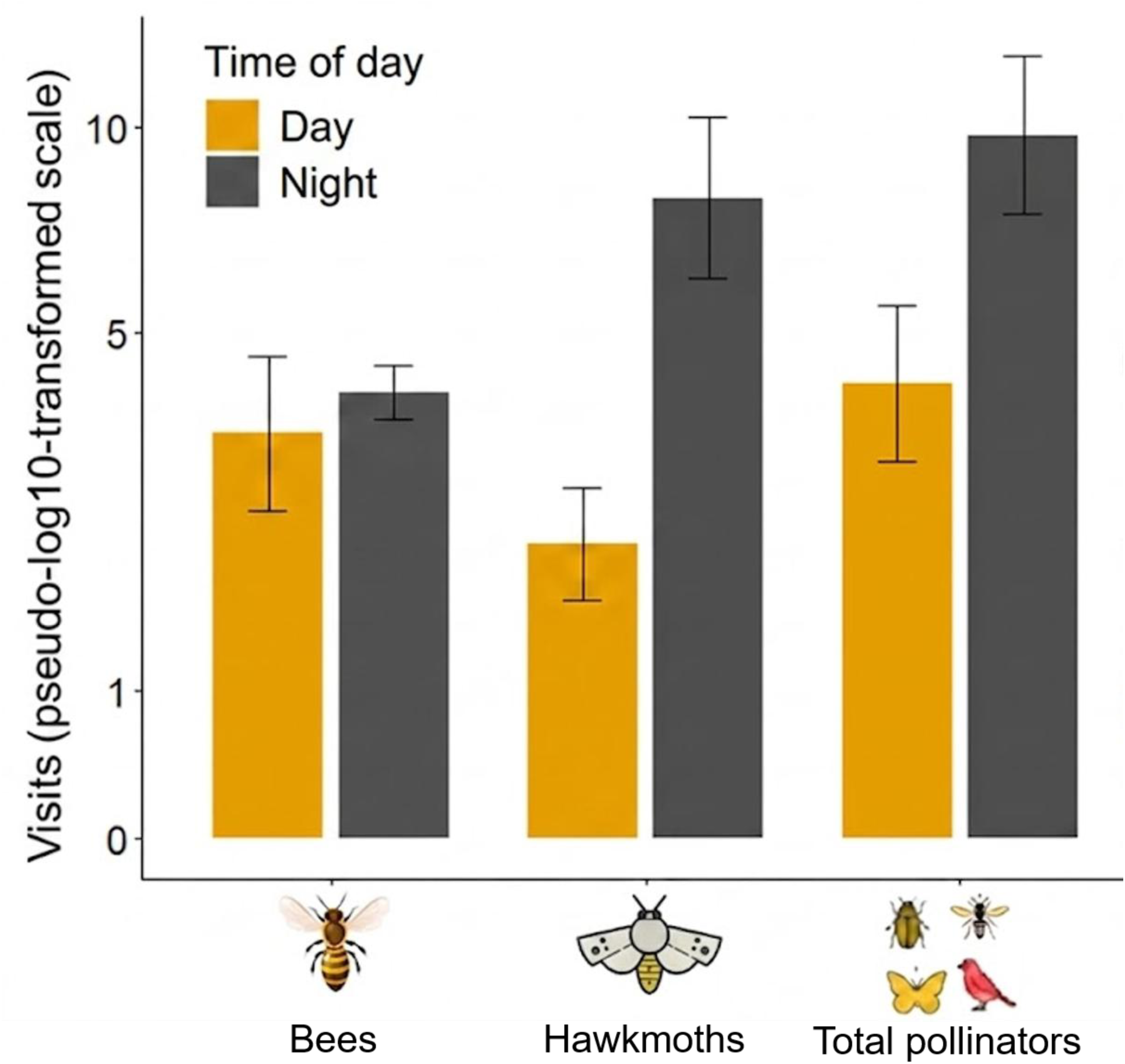
Temporal partitioning of floral visitors. While total visitation remains high across the diel cycle, the community is characterised by complete temporal partitioning. Bees dominate diurnal visits, whereas hawkmoths are the primary nocturnal visitors. Bars represent mean visits (± SE) on a pseudo-log10-transformed scale.

**Table 1:** Summary of generalised linear mixed effects models (GLMMs) evaluating the effects of farm management and landscape variables on floral visitor abundance. Results include parameter estimates, standard errors (SE), and z-values for fixed effects across six response categories.

| **Response variable** | **Explanatory variable** | **Estimate** | **Std. Error** | **z value** | **Pr(>\|z\|)** |
| --- | --- | --- | --- | --- | --- |
| Total flower visitors | (Intercept) | -2.176 | 0.536 | -4.062 | 0.000 |
|  | Farm management (organic) | 0.276 | 0.285 | 0.969 | 0.332 |
|  | Semi-Natural Habitats | -0.030 | 0.220 | -0.137 | 0.891 |
|  | Edge density | 0.405 | 0.220 | 1.841 | 0.066 |
|  | Diel period (night) | **-0.768** | **0.191** | **-4.029** | **<0.001** |
|  | Male or female tree | **4.110** | **0.433** | **9.495** | **<0.001** |
| Diurnal flower visitors | (Intercept) | 0.227 | 0.631 | 0.360 | 0.719 |
|  | Farm management (organic) | -1.026 | 0.743 | -1.381 | 0.167 |
|  | Semi-Natural Habitats | -0.121 | 0.592 | -0.205 | 0.838 |
|  | Edge density | 0.731 | 0.605 | 1.208 | 0.227 |
| Nocturnal pollinators | (Intercept) | -1.098 | 0.570 | -1.928 | 0.054 |
|  | Farm management (organic) | **1.233** | **0.626** | **1.970** | **0.049** |
|  | Semi-Natural Habitats | -0.195 | 0.496 | -0.394 | 0.693 |
|  | Edge density | -0.748 | 0.482 | -1.551 | 0.121 |
| Hawkmoths | (Intercept) | -1.499 | 0.634 | -2.364 | 0.018 |
|  | Farm management (organic) | **1.357** | **0.680** | **1.997** | **0.046** |
|  | Semi-Natural Habitats | -0.057 | 0.515 | -0.110 | 0.912 |
|  | Edge density | -0.802 | 0.513 | -1.562 | 0.118 |
| Other moths | (Intercept) | -2.189 | 0.667 | -3.280 | 0.001 |
|  | Farm management (organic) | -0.103 | 0.778 | -0.132 | 0.895 |
|  | Semi-Natural Habitats | 0.074 | 0.593 | 0.124 | 0.901 |
|  | Edge density | 0.519 | 0.578 | 0.897 | 0.369 |
| Bees | (Intercept) | -2.189 | 0.667 | -3.280 | 0.001 |
|  | Farm management (organic) | -0.103 | 0.778 | -0.132 | 0.895 |
|  | Semi-Natural Habitats | 0.074 | 0.593 | 0.124 | 0.901 |
|  | Edge density | 0.519 | 0.578 | 0.897 | 0.369 |

Total visitation was also strongly governed by male or female tree, with male trees receiving significantly higher visitation rates than female trees (z = 9.495, p < 0.001; Table 1).

The distance from a female tree to the nearest pollen donor (male or hermaphrodite *C. papaya*) exerted a significant effect on fruit set (*X*^2^ = 4.04, p = 0.045), generally shorter distances associated with higher fruit set than higher distances. In contrast, flower visitation rates by insect flower visitors remained unaffected by spatial proximity to male trees, showing no significant relationship with isolation distance (p>0.05).

### Experimental exclusion of pollinators Fruit set

We evaluated fruit set across five distinct pollination treatments: (1) open control for natural pollination, (2) hand-pollination to determine maximum potential yield, (3) total pollinator exclusion (closed) to isolate wind-only pollination, (4) day-exposure only, and (5) night-exposure only to evaluate temporal exclusion. Closed pollination yielded the lowest mean of 0.2 ± 0.13 fruit set per tree (Fig. 3a), while the mean number of papaya fruits set under open pollination was 4.1 ± 0.78 per tree, followed by day pollination (2.6 ± 0.65) and night pollination (2.0 ± 0.45) (Fig. 3b). Results from a generalised linear mixed effects model indicated that both day and night pollination treatments resulted in higher fruit set than closed pollination, with day pollination having similar effects than night pollination. Fruit set was significantly higher in flowers open to both diurnal and nocturnal pollinators than in those restricted to nocturnal pollination alone. However, fruit set did not differ significantly between the fully accessible flowers and the diurnal-only pollination treatment (Fig. 3d, Table S1).

**Figure 3:**
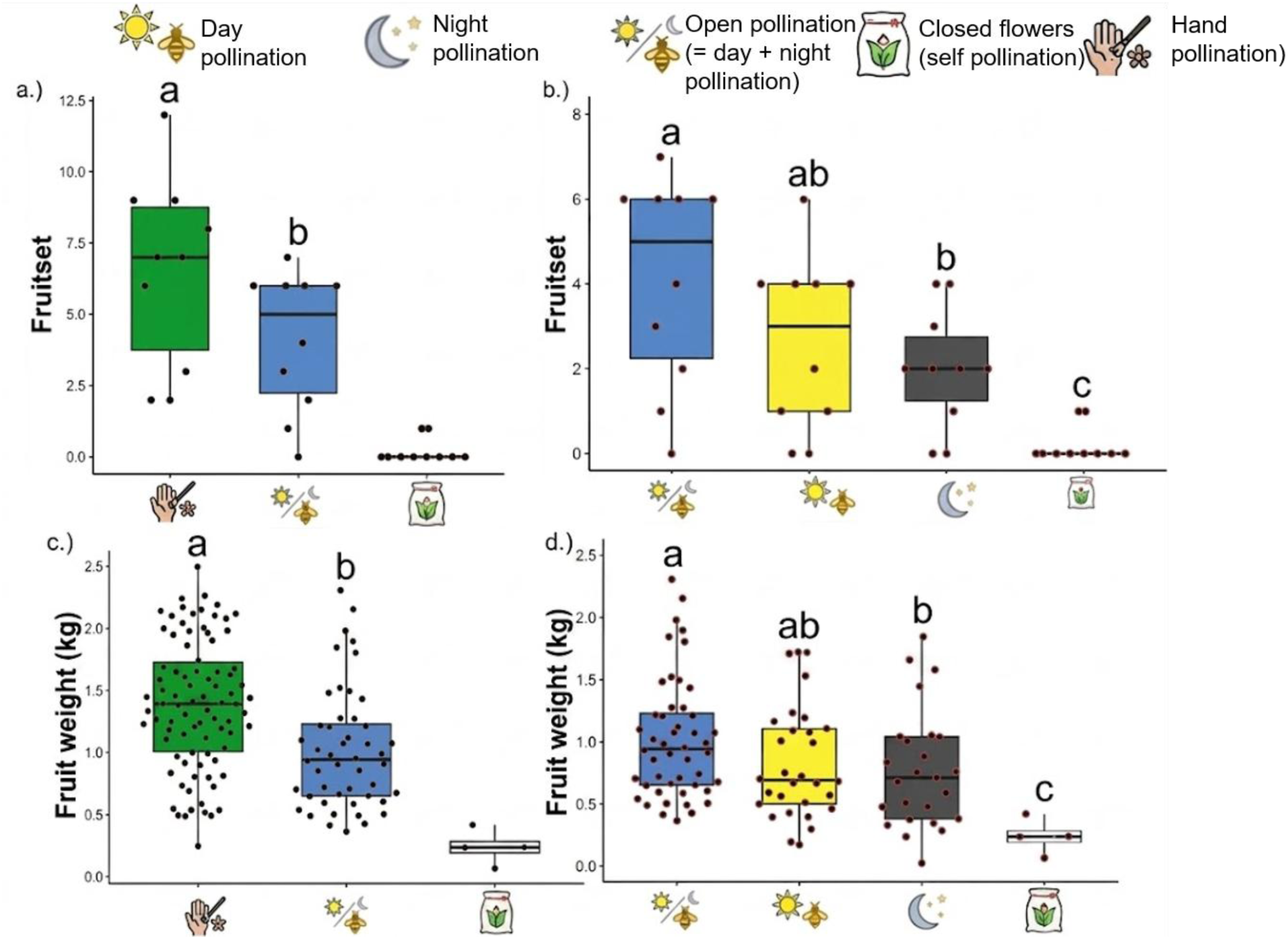
Reproductive success of *Carica papaya* across experimental pollination treatments. Impact of different pollination modes on fruit set at the site level (a, b) and individual fruit weight (c, d) are shown. **Left panels (a, c):** Comparison of fruit set per treatment per site (a) and individual fruit weight (c) under three baseline conditions: supplemental hand pollination (green n = 86 fruits), natural open pollination (blue n = 48 fruits), and closed flowers (white n = 4 fruits), which served as a control for autonomous/wind-mediated self-pollination. **Right panels (b, d):** Comparison of fruit set per experimental unit (b) and individual fruit weight (d) across diurnal and nocturnal pollination vectors: natural open pollination (blue, combining day and night, n = 48), day pollination only (yellow n = 30), night pollination only (grey n = 25), and closed flowers (white n = 4). Statistics for these treatments are shown in Table 2. *Note:* (*1*) *Data points (dots) represent different experimental units between metrics. For fruit set (panels a and b), each dot represents a single experimental tree/site (panels c and d), each dot represents an individual harvested fruit. Because a single branch often yielded multiple successfully set fruits, the total number of individual weight measurements is higher than the number of experimental trees. (2) The sample size for the ‘Closed’ (bagged) treatment is low (n* = 4*) due to extremely limited fruit development under total pollinator exclusion, reflecting a high reliance on biotic pollination*.

**Table 2:** Pollination treatment effects on papaya reproductive success. Results of Generalised linear mixed effects models (GLMM) for fruit set and linear mixed-effects models (LMM) for fruit weight and total production are shown. All models include Site ID as a random effect and use Open pollination (combined day and night) as the reference level for the intercept.

| **Response Variable** | **Explanatory variable** | **Estimate** | **Std. Error** | **z/t value** | **P value** |
| --- | --- | --- | --- | --- | --- |
| Fruit set | Intercept | **1.344** | **0.203** | **6.626** | **<0.001** |
|  | Day pollination treatment | -0.456 | 0.249 | -1.832 | 0.067 |
|  | Night pollination treatment | **-0.718** | **0.27** | **-2.654** | **0.008** |
|  | Closed flowers (wind) treatment | **-3.02** | **0.718** | **-4.207** | **<0.001** |
| Fruit weight (g) | Intercept | **1030.78** | **76.05** | **13.554** | **<0.001** |
|  | Day pollination treatment | -202.46 | 107 | -1.892 | 0.061 |
|  | Night pollination treatment | **-265.59** | **113.76** | **-2.335** | **0.022** |
|  | Closed flowers (wind) treatment | **-812.74** | **237.67** | **-3.42** | **<0.001** |
| Production | Intercept | **4.12** | **0.51** | **8.06** | **<0.001** |
|  | Day pollination treatment | **-1.90** | **0.67** | **-2.838** | **0.009** |
|  | Night pollination treatment | **-2.58** | **0.67** | **-3.848** | **<0.001** |
|  | Closed flowers (wind) treatment | **-4.10** | **0.67** | **-6.118** | **<0.001** |

### Fruit weight and yield

Analysis of papaya fruit weight under different pollinator exclusion treatments showed significant differences in mean weight. The comparison between open and night pollination demonstrated a statistically significant difference in fruit weight of 266 grams (Fig 3d).

We examined the effect of diurnal and nocturnal pollination treatments on total papaya production (yield, measured as fruit set x average weight; Fig. 4). To assess whether diurnal and nocturnal pollinators contribute to yield in an additive or synergistic manner, we compared the yields of the single-period treatments (diurnal-only and nocturnal-only) to the unrestricted open-pollination treatment. The cumulative yield of the separate day and night treatments closely approximated the total yield achieved under open-pollination conditions. The open-pollination yield did not significantly differ from the predicted additive sum of the two single-period treatments (t = 0.402, p = 0.691).

**Figure 4:**
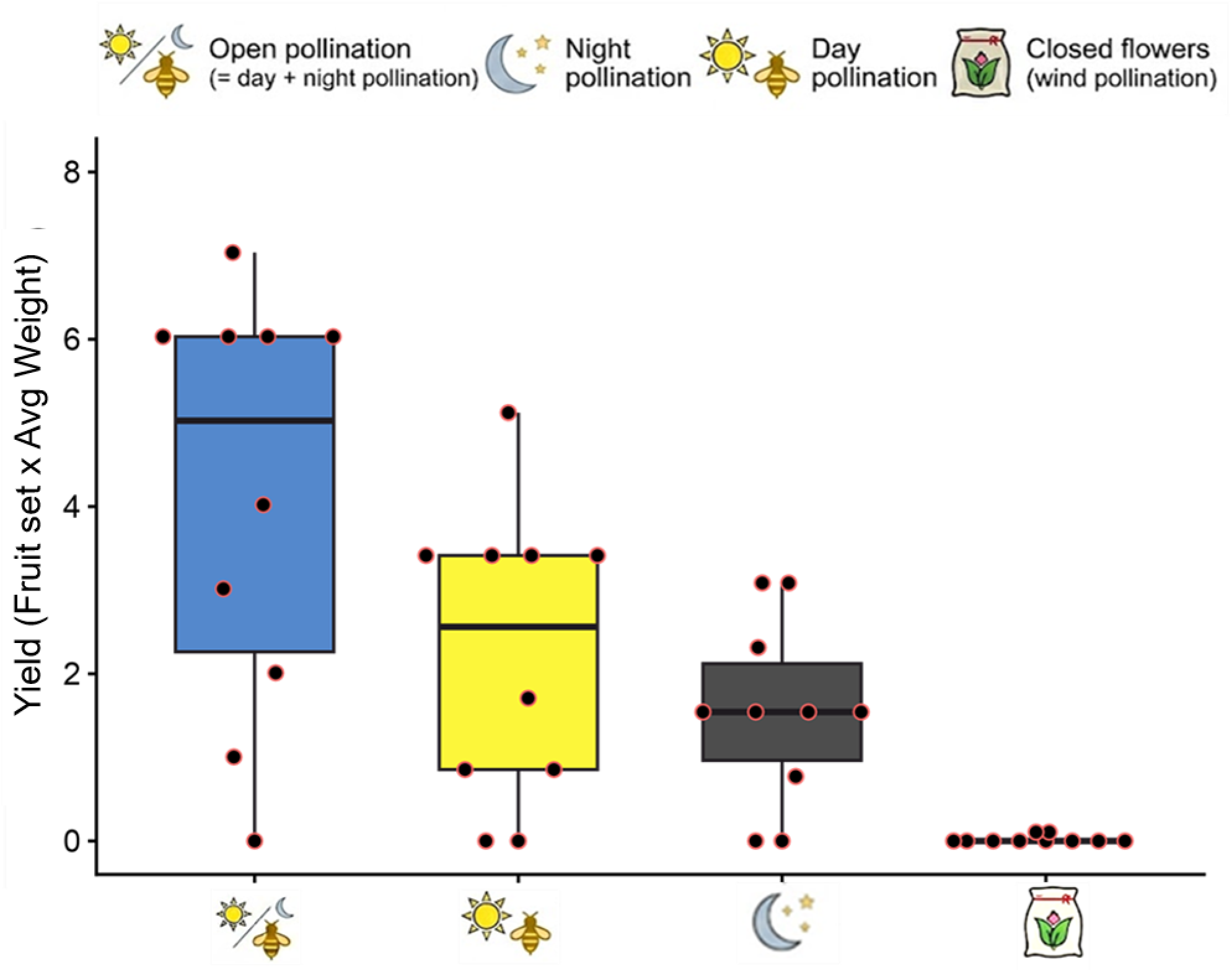
Cumulative papaya production across pollination treatments. Total production (calculated as fruit set x average fruit weight) is significantly higher under open pollination (blue) than under day (yellow) or night (grey) pollination.

Pairwise comparisons revealed that open pollination significantly enhanced fruit set compared to night-only exclusion, resulting in a 2.05-fold increase in reproductive success (z = 2.654, p = 0.008; Table S1). We found no statistically significant differences in fruit set between open and day-pollinated flowers (p = 0.067) nor between night and day-pollinated flowers (p = 0.374; Table S1).

### Landscape and local management effects on pollinator activity and yield components

To understand how landscape context and local management shape pollinator communities, we evaluated the relative influence of landscape composition (SNH), configuration (edge density), and farm-level practices on the abundance and activity of papaya flower visitors. Our models revealed that when flower visitors were analysed as a single, combined pool, overall insect visitation was largely unaffected by local farm management, showing only a marginally positive trend with landscape configuration (z = 1.841, p = 0.066; Table 1). Instead, overall visitation patterns were primarily driven by biological and temporal factors, such as male or female tree and sampling time.

Evidence for guild-specific sensitivity to management practices emerged when disaggregating the visitation data by temporal niche. While the total visitor pool showed no significant response to management practices (Table 1), analysing pollinators by guild reveals a more complex pattern of response to these environmental drivers. Nocturnal pollinators demonstrated a statistically significant positive response to organic management (z = 1.970, p = 0.049; Fig. 5a), a trend particularly pronounced in hawkmoths (z = 1.997, p = 0.046; Table 1). In contrast, diurnal flower visitors, including bees, did not show significant responses to local farm management, nor were they significantly influenced by landscape composition or configuration (p > 0.05 for all explanatory variables; Table 1).

**Figure 5:**
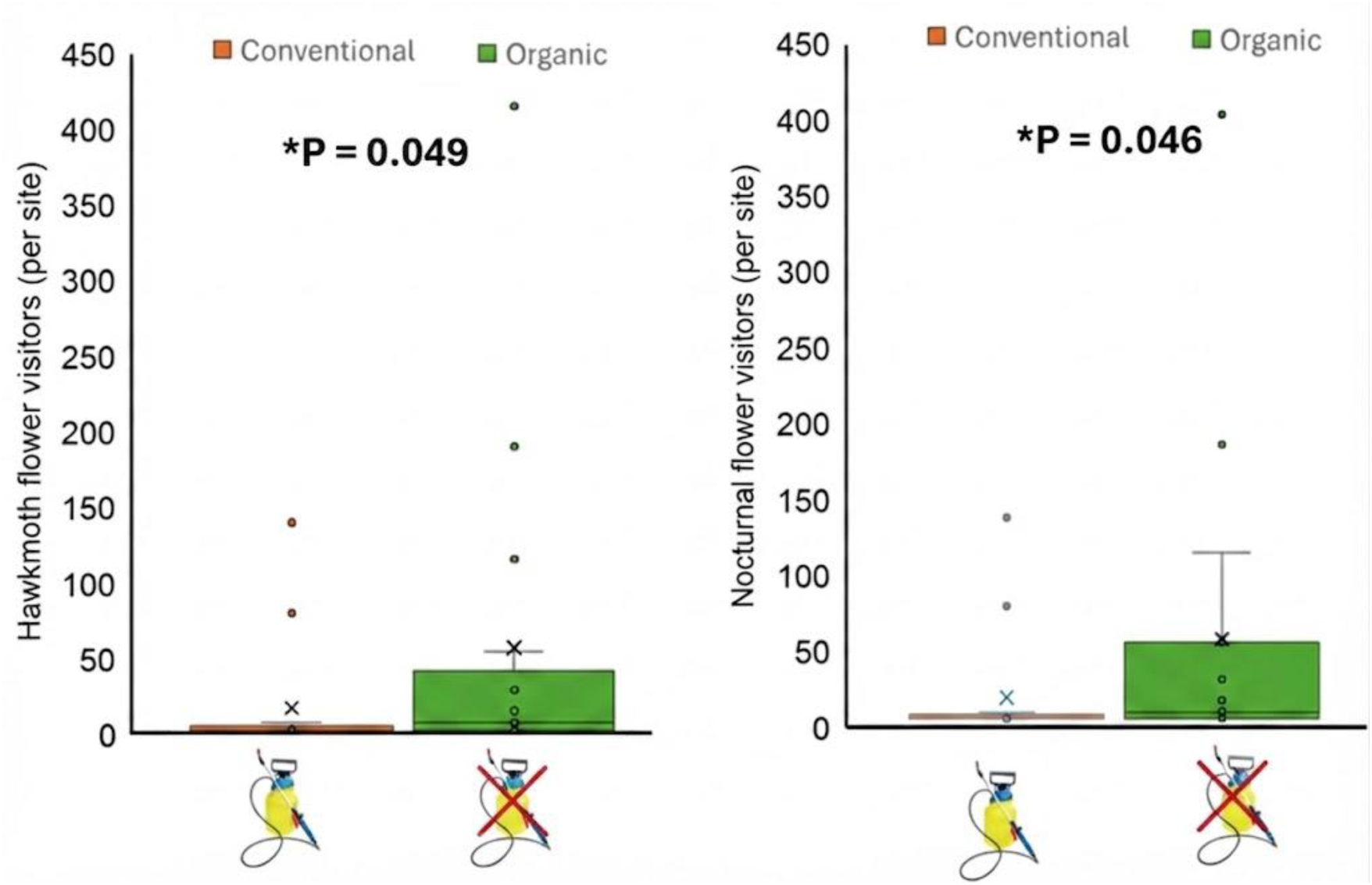
Impact of local farm management on the abundance of nocturnal floral visitors. Box-and-whisker plots illustrating the response of (a) hawkmoths (Sphingidae) and (b) total nocturnal flower visitors to conventional versus organic farming practices. Abundance is expressed as the total number of visits recorded per site (summed across all 10-minute observations per tree across all trees at that site). Conventional sites (N=6) are represented by an insecticide sprayer icon, signifying the use of synthetic chemical inputs, while organic sites (N=10) are represented by a crossed-out sprayer icon, signifying the absence of synthetic chemical inputs. *One farm was excluded from analyses due to insufficient data regarding farm management practices.

We found no significant relationship between pollinator activity, categorized by diurnal and nocturnal visitors, or farm management practices and the total fruit set (Table S2). The Negative Binomial GLM indicated that diurnal visitor activity (z = –1.071, p = 0.284), nocturnal visit activity (z = 0.280, p = 0.779), and farm management (organic versus conventional; z = –1.022, p = 0.307) were not significant predictors of total fruits set in this study. Similarly, our analysis of fruit quality revealed no significant relationship between pollinator activity or management practices and average fruit weight (Table S2). The model indicated that diurnal visitor activity (z = –0.250, p = 0.803), nocturnal visit activity (z = –0.192, p = 0.848), and farm management (organic vs. conventional; z = –0.125, p = 0.900) were not significant predictors of fruit weight. Further, neither diurnal nor nocturnal visitor abundance, nor farm management practices, significantly influenced crop yield Table S2. The analysis showed no significant association with diurnal visitor activity (z = –1.071, p = 0.284), nocturnal visitor activity (z = 0.280, p = 0.779), or Farm management (z = –1.022, p=0.307).

## Discussion

Our study shows that papaya pollination is sustained by complementary pollination services provided by diurnal and nocturnal insects, with strong temporal niche partitioning. While bees dominated daytime visitation, nocturnal hawkmoths emerged as highly efficient flower visitors. Nocturnal pollinators were more sensitive to local management, with organic farming enhancing hawkmoth abundance. The approximately 36.9% yield gap between natural and supplemental hand pollination highlights the potential of pollinator management for sustainable tropical papaya production.

### Pollen limitation and the yield gap

We found that papaya production strongly depends on pollination services and that crops in the study region face significant pollination limitations. Large deficits in fruit set (36.9%) and fruit weight (32.40%) when comparing trees with supplemental hand pollination to controls (open flowers) indicate that papaya production is strongly limited by the amount of pollen received by the flowers. These deficits are higher than the global average of 20%-25% for tropical crops [21]. This substantial gap (36.9%) suggests that pollinator communities in the study region do not provide pollination services at a level sufficient to reach full reproductive potential. Importantly, this gap likely reflects both a lack of pollinator abundance and a decline in the quality of pollination services, suggesting that the current pollinator community is failing to provide the high-quality pollen transfer required for optimal fruit set and weight.

One might question whether this severe pollination gap is a consequence of the crop being grown outside its native Neotropical range, where specialised co-evolved pollination vectors might be absent. However, pronounced pollen limitation is a globally documented characteristic of *Carica papaya*, occurring extensively within both its native range in the Americas and its introduced paleotropical range [22]. Because female papaya flowers are entirely rewardless, they visually mimic male flowers to deceive visitors seeking nectar-rich rewards [23]. As a result, low visitation rates and substantial pollen deficits limit fruit set globally, irrespective of geographic origin. Furthermore, key functional pollinator guilds are not missing in Africa; rather, highly mobile paleotropical hawkmoth communities, including native East African species such as *Hippotion celerio*, *Nephele comma*, and *Agrius convolvuli,* perfectly mirror the ecological niche filled by Neotropical sphingids, executing over 95% of legitimate cross-pollination events when local farm management and nearby natural habitats provide adequate resources [24].

The 36.9% drop in fruit set shows that the pollen delivered by natural pollinators is often insufficient or of poor quality. Since fruit weight in papaya is often linked to seed set, this weight loss likely means that not all ovules are fertilised. Given the additive services of both diurnal and nocturnal guilds (Fig. 3), the significant limitation observed here suggests a decline in efficiency or a lack of habitat connectivity for key groups, such as hawkmoths or bees. However, this pollination gap likely stems from a combination of multiple proximate and ultimate drivers rather than isolation alone. Proximately, intensive agricultural practices impose severe localised disturbances, including pesticide drift and vegetation clearing, which directly deplete overall pollinator abundance and disrupt foraging behaviour [25]. Ultimately, the degradation of surrounding landscapes leads to a critical loss of essential larval host plants and nesting substrates within the regional species pool, preventing specialised nocturnal and diurnal vectors from establishing viable populations [26]. Thus, the observed decline in pollination services is likely a multifaceted consequence of habitat fragmentation, reduced floral resource diversity, and management-induced disturbances that collectively filter out highly efficient, specialised pollinators from the agricultural matrix.

### Complementary roles of diurnal and nocturnal pollinators

Our observation of 2,056 floral visitors highlights a community dominated by two primary functional groups: bees and hawkmoths, which together accounted for nearly 90% of all visits. This finding suggests that papaya effectively utilises two distinct temporal niches to ensure reproductive success, a pattern also observed in other tropical plants to secure pollen transfer when pollinator availability fluctuates [13,27]. While this dual-niche utilisation may represent an adaptive bet-hedging strategy to mitigate the risks of pollen limitation across variable environments [28], it could also simply reflect a generalised floral phenotype that successfully recruits diverse visitor guilds, without necessarily reflecting a specific evolutionary history of temporal partitioning [29].

The clear temporal niche partitioning observed may reduce interspecific competition for floral resources and ensure that the plant receives adequate pollination services. Our data show that while diurnal and nocturnal treatments provided statistically similar fruit set individually (Table 2; Fig. 3), the ‘Open’ treatment (both guilds present) resulted in a significantly higher fruit set. This pattern suggests that these pollination services are cumulative rather than redundant; if the guilds were functionally redundant, the ‘Open’ treatment would not significantly outperform the single-period treatments. Furthermore, the increase in the ‘Open’ treatment indicates that the services provided by day and night guilds are additive, providing continuous pollination coverage throughout the 24-hour cycle. While our results demonstrate an additive pattern, the potential for synergistic effects, such as the delivery of pollen loads with varying quality or compatibility that might enhance fruit development beyond the sum of their individual contributions, remains a hypothesis for future investigation.

While bees are often the most visible pollinators in agricultural systems [30], our data suggests that for papaya, the nocturnal shift, driven by hawkmoths, is a vital and nearly equal partner in the production cycle. Although diurnal pollination provides a slightly larger share of the total yield, our findings confirm that removing either the diurnal or nocturnal component would significantly impact overall productivity [31,32].

### Pollinator attraction of male and female trees

Our findings demonstrate that broad-scale landscape metrics, such as the proportion of surrounding landscape composition, fail to explain local variations in papaya reproductive success. Instead, fruit set is governed by localised spatial configuration, specifically the isolation distance from pollen donors. We observed a highly significant pollinator bias toward male trees (z = 9.495, p < 0.001). Because male flowers in dioecious papaya species offer both pollen and nectar while female flowers offer no rewards, pollinators are naturally drawn to these reward-heavy males. We refer to the lack of nectar rewards in female flowers as ‘mimicry by default’ [33]. This term is used here to describe a resource-based deceptive strategy, wherein female flowers gain visitation by resembling the rewarding male flowers, rather than a specific morphological adaptation resulting from a co-evolutionary mimicry trait [33, 34]. It is important to clarify that this ‘mimicry’ is functional and context-dependent, stemming from the floral display’s lack of reward rather than the development of specialised visual or olfactory signals designed to deceive pollinators [33].

While hawkmoths are highly sensitive to floral volatiles [35,36], current evidence suggests limited divergence in scent profiles between male and female papaya flowers [36,37]. This lack of distinct olfactory signalling may further facilitate this ‘default’ mimicry, as pollinators, unable to distinguish between sexes through scent, are more likely to visit female flowers by mistake while foraging for the high-energy nectar rewards emitted by the nearby male population [36]. While scent uniformity is a consistent trait reported in previous research [36,37], it remains unknown whether this is a universal characteristic across all *Carica* varieties or if it is specific to the populations within our study region. Importantly, as fast-learning foragers, hawkmoths might theoretically learn to associate specific floral displays with a lack of reward, potentially avoiding female flowers over time. However, we hypothesize that the higher abundance of pollinators on organic farms may actually slow this learning process; in these resource-rich landscapes, a higher frequency of male flowers and a denser, more diverse floral community may act as a source of constant distraction, reducing the pollinators’ ability to discriminate between rewarding and non-rewarding individuals.

This persistent visitation, driven by a higher density of foraging events, may therefore bridge the yield gap between organic and conventional systems by ensuring female flowers receive consistent pollination services despite their deceptive nature.

This disparity in attraction makes the spatial arrangement of male and female trees important. If male trees are too far away, these accidental visits to female trees may drop below the threshold required for optimal pollination. Our empirical data strongly support this mechanism: we observed a significant decline in fruit set as the distance to the nearest male or hermaphrodite tree increased, even though insect visitation rates remained stable across the same spatial gradient. However, a marginal positive correlation with landscape configuration (z = 1.841, p = 0.066) suggests that complex landscapes, which increase the interface between different tree types, may facilitate pollinator movement between male and female plants and help overcome this attraction bias [38]. This potential driver of connectivity highlights that edge density likely modulates the efficiency of inter-tree foraging; to ensure these parameter estimates were robust and to account for model selection uncertainty, we performed full model averaging [30,59], which consistently identified landscape configuration as a relevant, albeit marginal, contributor to observed visitation patterns.

The high visitation rate of hawkmoths can be attributed to their high-energy foraging behaviour [39]. Their large body sizes and hover-feeding style result in effective contact with the reproductive organs of papaya flowers [24]. Furthermore, papaya flowers often exhibit nocturnal anthesis, meaning pollen is most viable and nectar most abundant when hawkmoths are active. This aligns with previous research noting that while many tropical fruit crops are visited by bees, they rely heavily on nocturnal insects to achieve their full reproductive potential [40].

### Landscape and local management effects and the resilience of pollinators

Landscape and local management effects and the resilience of pollinators We found that local-scale ecological intensification through organic management influences pollinator communities more strongly than landscape-scale factors. The Taita Hills contain a complex, heterogeneous mix of semi-natural habitats that likely buffers the system, masking potential landscape-scale effects of degradation [4]. In these agricultural mosaics dominated by semi natural habitats, ecological differences between patches are low, meaning landscape-level impacts may only appear if the area undergoes severe structural simplification, such as large-scale conversion into homogeneous monocultures [41,42]. Furthermore, this lack of landscape-scale response could result from a spatial mismatch between our metrics and the scales at which different pollinator groups operate [43]. While smaller diurnal bees forage locally, highly mobile hawkmoths integrate resources across a broader landscape, effectively smoothing out variations in habitat composition.

Our finding that the total pollinator community shows no response to management while specific nocturnal guilds thrive provides important insights. By aggregating all visitors into a single metric, community-wide analysis likely obscures the specialised responses of hawkmoths, which are more sensitive to agricultural inputs than the more diverse diurnal visitor pool. This “masking effect” suggests that traditional biodiversity metrics may be insufficient for evaluating conservation in tropical systems where different functional groups have distinct life-history requirements. Consequently, guild-specific analysis is a methodological necessity. This approach is essential for identifying how management practices, such as the preservation of understory larval host plants, facilitate the survival of key nocturnal vectors, even when those benefits are not visible in the total pollinator abundance.

While organic management significantly increased the abundance of nocturnal pollinators, particularly hawkmoths (Fig. 5), this did not translate into statistically significant improvements in papaya fruit set or fruit weight, or cumulative yield compared to conventional sites. This discrepancy highlights a critical distinction between regional pollinator supply and the actualised delivery of crop services. Three mechanisms may explain this: first, the relationship between pollinator abundance and reproductive output may reach a saturation threshold, where the benefits of increased nocturnal visitors in organic fields are offset by other physical limitations, such as soil nutrients, water availability, or maternal investment capacity. Second, highly mobile diurnal visitors, which did not differ between management types, may already provide a baseline of pollination sufficient to meet the crop’s maximum reproductive potential, rendering the additional nocturnal visits functionally redundant for immediate yield.

Third, the complex, resource-rich landscape matrix of the Taita Hills likely sustains a high regional baseline of pollination services that masks localised management contrasts at the farm scale. Regarding the quality mismatch hypothesis, our current dataset does not include direct measures of pollen load or stigma pollen receipt; consequently, we addressed these potential drivers by performing full model averaging to account for model selection uncertainty and to ensure that our conclusions regarding the limited effect of visitor abundance on reproductive output remain robust across the candidate model set [30,59].

Furthermore, the persistence of a 36.9% yield gap despite higher pollinator numbers on organic farms may reflect a fundamental mismatch between pollinator quantity and pollination quality. Because female papaya flowers are rewardless and rely on food deception, hawkmoths—which are highly cognitive, fast-learning foragers—may rapidly learn to avoid these flowers in favour of rewarding male or hermaphrodite flowers [33]. This behavioural avoidance, combined with spatial mismatches in pollen-donor distribution, suggests that localised farm-level increases in hawkmoth abundance do not necessarily translate into effective, high-quality pollen transfer. Future research must prioritize investigating the interplay between hawkmoth learning behaviour, floral reward structures, and the efficacy of pollen deposition to determine if these cognitive factors are the primary constraints limiting the translation of increased pollinator abundance into higher crop yields.

However, the lack of an immediate yield difference should not obscure the conservation value of local management. The higher abundance of efficient nocturnal vectors on organic farms suggests that these practices build essential ecological resilience. By eliminating synthetic inputs and preserving diverse understory larval host plants such as *Ipomoea* spp., *Triumfetta rhomboidea*, and *Eleusine indica* [44], organic farming supports the full life cycle of the local sphingid community. Consequently, organic management acts as a vital ‘ecological safety net.’ While it may not drive immediate yield spikes under optimal conditions, sustaining these robust, dual-niche pollinator communities is highly likely to stabilise agricultural production against environmental fluctuations and long-term climate destabilisation [14].

Bees, conversely, operate at smaller spatial scales and may be primarily constrained by the immediate availability of nesting substrates in adjacent patches rather than broad-scale landscape configurations [45]. Our findings demonstrate that by boosting the hawkmoth community, organic management potentially enhances nocturnal pollination services [46].

This study underscores that pollinator conservation in tropical agriculture must be broadened to include nocturnal taxa. The reliance on hawkmoths, a group often overlooked compared with honeybees, means that practices such as maintaining specific larval host plants are important for crop security [47]. Unlike generalist bees, hawkmoths require specific vegetation for their caterpillars, making the conservation of non-crop host plants a critical component of integrated management. Furthermore, the role of organic management in supporting hawkmoths provides a clear economic incentive for pollinator-friendly farming. By fostering a diverse insect community that operates across the full daily cycle, farmers can move toward a more self-sustaining ecosystem. Protecting complementary diurnal and nocturnal pollination services will be essential for securing global food production [48].

## Conclusion

Our results show that nocturnal pollination is an important driver of successful papaya reproduction in tropical areas, yet it remains the most vulnerable component of the pollinator community under intensive farming. From a farming perspective, losing both the quantity and quality of papaya fruit means farmers earn far less income than they could. These findings show there is a strong likelihood of improving yields by focusing on ecological solutions. If we can close the 36.9% yield gap through improved habitats or farming practices, smallholder farmers could see significant gains in food security and income without increasing external inputs. Protecting these 24-hour networks is essential for the long-term health and economic success of tropical agriculture.

Importantly, the lack of a significant impact of landscape-scale factors suggests that the biological signal of the surrounding mosaic is secondary to the immediate benefits of local farm management. This provides a powerful incentive for smallholder farmers: local transitions to organic practices, such as avoiding synthetic chemicals and maintaining diverse understory larval host plants, can significantly enhance pollination services and yields regardless of the surrounding landscape context.

## Materials and Methods

### Study region

The study was conducted in the Taita Hills of Southeast Kenya (Fig. 6), a region characterised by a sharp climatic divide between the highlands and the surrounding lowlands, from January 2024 to December 2025. The hills act as a natural barrier, intercepting moist winds from the Indian Ocean and resulting in persistent mist and cloud cover at higher altitudes. This unique environment keeps the local forests wet all year-round, with annual precipitation averaging between 1,332 mm and 1,910 mm. While the region traditionally follows two distinct rainy seasons, a long one from March to May and a shorter one from November to December, weather patterns have recently become increasingly unpredictable, likely a consequence of global climate change. In sharp contrast, the adjacent lowlands are arid, receiving as little as 250 mm of rain per year.

**Figure 6:**
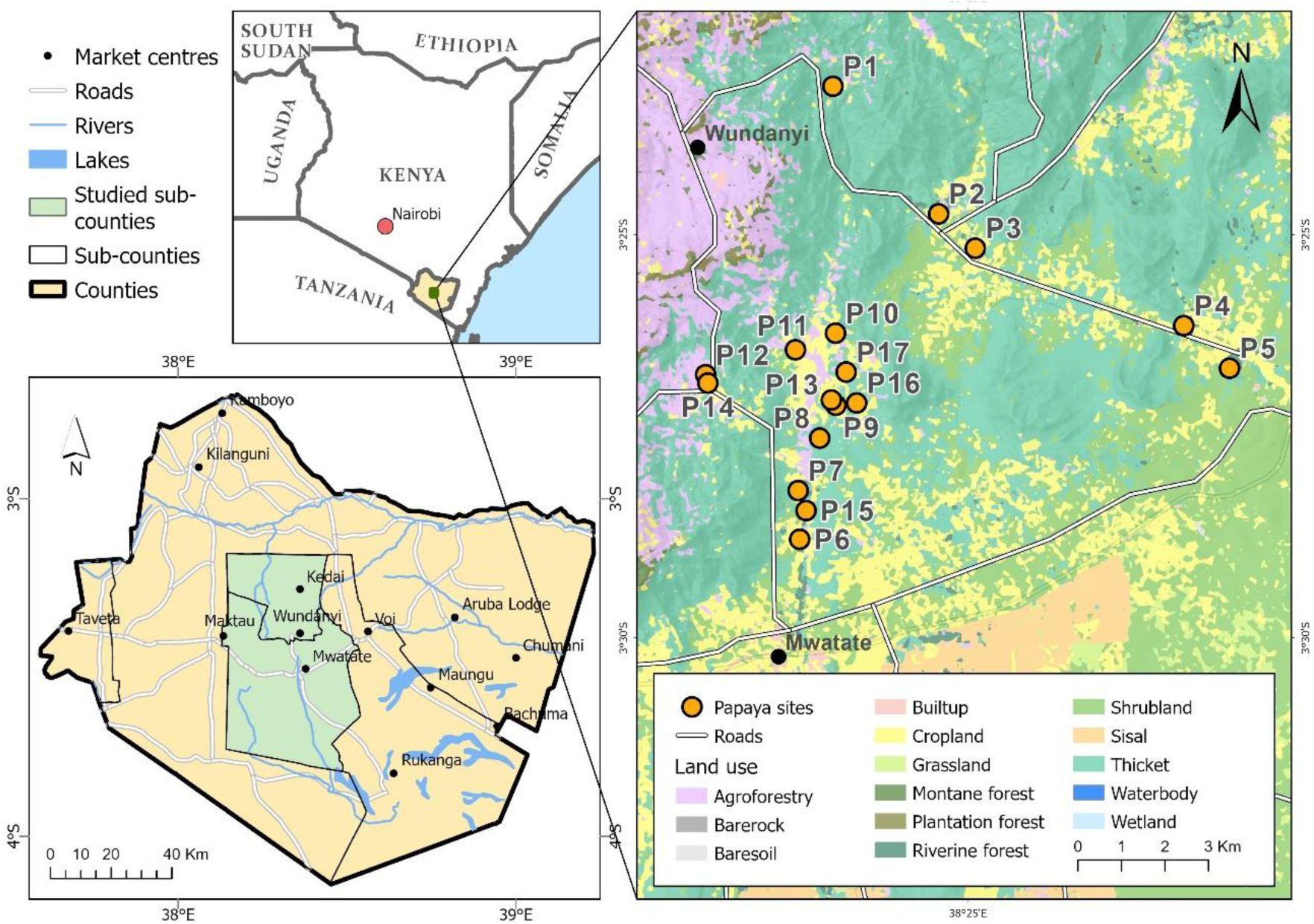
A map of the study area within the Taita Hills (inset map of Kenya).

The region’s distinct climate supports a high degree of biodiversity, particularly among invertebrates. The Taita Hills are home to a wide diversity of flies and moths, including three endemic butterfly species and a high diversity of bees [49]. Beekeeping for honey production (mostly honeybees but also stingless bees) is a common practice throughout the hills.

Agriculture in the area is characterised by smallholder farms that primarily produce food for personal consumption or small-scale local trade. These farms are mostly organic, as they are typically managed with low-intensity methods that involve minimal intervention beyond planting, weeding, and basic irrigation. While maize and beans are the primary crops, grown twice a year during the rainy seasons, the farms are notably diverse. Most farmers cultivate a wide range of additional crops in smaller quantities, including staples and fruits such as papaya, pigeon pea, cassava, banana, chilli, mango, avocado, sweet potato, and guava.

### Study design

To address our objectives, we selected 17 *Carica papaya* farms over a two-year period, combining continuous observations of nocturnal and diurnal floral visitors with experimental pollination-exclusion treatments. The study sites were positioned along a landscape-level land-use intensification gradient, with proportional cropland coverage ranging from less than 10% to greater than 90% within a 500 m radius of each farm. To ensure that our observations were not confounded by altitudinal variations, all fields were selected within a narrow elevation band between 650 m and 1137 m above sea level.

This design integrated both surrounding landscape context and local crop management practices to assess how these dual scales of intensification shape pollination services and papaya yield parameters. To evaluate local management, we assessed the history of agricultural practices at each site through a combination of farm record reviews and face-to-face interviews with farmers. We categorised the 17 study sites based on their agrochemical management. Six farms were classified as conventional, as they utilised synthetic inputs, while 10 were classified as organic, utilising biopesticides, biofertilizers, or no inputs at all. One farm remained uncategorized; although we were granted research access, the farmer was unavailable for an interview to confirm their specific management practices. Ultimately, this integrated framework allowed us to directly compare the activity, community composition, and functional contributions of diurnal and nocturnal pollinators across varying local and landscape-level environmental conditions.

### Flower visitor observations

Flower-visitor observations were conducted during two flowering seasons (March–April 2024 and March–April 2025) using a repeated-sampling-session protocol. Each sampling unit consisted of 10 minutes of visual observation, immediately followed by 5 minutes of specimen collection; this cycle was repeated continuously during each session. During observation intervals, two observers remained close to focal flowers, minimising movement and noise to avoid disturbing visitors, and recorded for each visit the time, whether the flower was male or female, number of visits, visitor behaviour (e.g., landing, probing, nectar or pollen foraging) and a field-level taxonomic identification (e.g., bee, moth, fly).

Diurnal observations were conducted between 10:00 and 16:00 under stable weather conditions, characterised by sunny to moderately cloudy skies and low wind. Nocturnal sampling was scheduled for 18:30–20:00, targeting the peak visitation window identified during preliminary observations at 18:00 and 06:00, which indicated that nocturnal pollinator activity began shortly after sunset (∼18:40) and then declined over the subsequent hour (∼20:00). Both diurnal and nocturnal sampling periods followed identical protocols, consisting of a 10-minute visual observation cycle followed by 5 minutes of sweep-netting. To ensure consistency and minimise disturbance, infrared headlamps were utilised during all nocturnal sessions.

We identified primary pollinators with distinct features, such as honeybees (*Apis mellifera*) and most hawkmoths (e.g., *Agrius convolvuli*), to the species level in the field. For other flower visitors that could not be identified on site, we collected voucher samples using sweep nets, euthanised them with ethyl acetate, and took them to the field station for later identification in the lab. In the laboratory, we examined the specimens under a stereomicroscope to assess diagnostic features, including wing venation and leg shape. We used taxonomic keys [e.g. 32] to identify them to genus or species when possible, or to morphospecies when needed. The collected specimens are stored at the University of Embu’s Entomology Lab.

### Pollination treatments

Pollination experiments comprised five treatments: open (natural pollination), closed (wind pollination), hand pollination (pollen limitation), day-only pollination, and night-only pollination. Flowers were labelled, and treatments were applied to newly emerged female flowers on tagged trees during late January to mid-April 2024 and 2025. Treatments were assigned in alternating order, when possible, but actual assignment depended on flower availability and anthesis timing (anthesis ≈ 3 days). Labelled flowers were checked regularly and excluded if they died before or during anthesis.

Open (natural pollination) treatment: A focal flower was marked and left entirely exposed to natural conditions; no manipulation was performed.

Closed (wind pollination) treatment: A gauze bag (mesh opening 0.3–0.5 mm) was fitted around each flower while still closed so that no insect visitor could access the flower once it opened, while allowing air movement and potential wind-borne pollen. Bags remained in place throughout anthesis and were removed as soon as the flower visibly wilted to avoid mechanically interfering with early fruit development.

Hand pollination treatment: Female flowers at anthesis were hand-pollinated using fresh pollen from three different male trees. For each hand pollination event, three freshly opened male flowers (one at a time, from different male trees) were collected, their petals removed, and the anthers gently rubbed across the female stigma for ∼3 seconds per male flower. Male flowers showing discolouration or other signs of senescence were excluded. The stigma was photographed before and after pollination for documentation.

Day-only and night-only pollination treatments (temporal exclusion): For both treatments, a gauze bag (0.3–0.5 mm openings) was attached around each flower while still closed. Bags were then managed on a daily schedule based on preliminary pollinator-activity observations and local sunrise/sunset times (sunrise ≈ 06:30, sunset ≈ 18:40): bags were removed every morning between 07:00 and 07:30 and reattached every evening between 17:45 and 18:15. During the evening reattachment period, we occasionally observed low numbers of hawkmoths scouting or hovering near focal trees; however, these sightings were infrequent and did not reach the high visitation frequencies recorded during the formal nocturnal sampling window (19:00–20:00) when anthesis peaked. For the day-only treatment, the bag was removed each morning and reattached each evening, so the flower was uncovered during the day and inaccessible at night. For the night-only treatment, the bag remained off overnight, was reattached in the morning, and removed in the evening, so the flower was exposed only during nocturnal hours. This schedule targeted observed peaks of diurnal and nocturnal visitation while minimising unintended exposure.

To evaluate the spatial drivers of reproductive success, we monitored all female *C. papaya* individuals bearing flowers accessible from ground level. Each target female tree was assigned a unique identifier, and we recorded its Euclidean distance to the nearest sexually mature, flowering pollen donor (either male or hermaphrodite *C. papaya*).

While we sampled 17 distinct sites for flower visitors and pollination over the two-year study, the distribution of the experimental pollination treatments varied by year. In 2024, basic fruit set experiments (comprising open and closed treatments) were conducted across 13 sites, with one representative site receiving the full suite of all five experimental pollination treatments. In 2025, we expanded the scope of the experimental setup, successfully conducting all five pollination treatments across 13 study sites. For all treatment implementations, we standardised our sampling by targeting five trees per site and tagging five flowers per treatment per tree. Throughout anthesis and early fruit development, these labelled flowers were checked frequently to record their fate.

Flowers that died before or during anthesis were excluded from analysis. Harvests were conducted from mid-June to the end of October 2024 and repeated in the same months in 2025. After harvest, fruits were stored at ambient room temperature until fully ripe; fruit weights were recorded using a Digital Kitchen Scale with a precision of ±1g.

### Farm management measurement

To evaluate the impact of agricultural intensification on pollination services, study sites were classified into two distinct management categories, organic and conventional, using longitudinal data obtained through pre-tested, semi-structured interviews with papaya farmers. These interviews were designed to capture a three-year history of input use, ensuring that the observed ecological effects represented sustained management regimes rather than transient seasonal variations.

Farms were classified as organic if they relied exclusively on ecological intensification techniques, such as applying farmyard manure and compost, and using botanical or biological pesticides. These organic systems were further characterised by maintaining a diverse understory and not using synthetic agrochemicals. In contrast, conventional sites were characterised by the routine application of synthetic agrochemicals, specifically inorganic fertilisers such as NPK or CAN, and broad-spectrum synthetic pesticides, including neonicotinoids and pyrethroids, for pest and disease control.

To isolate management effects from environmental effects, sixteen representative *C. papaya* orchards were selected across the Taita Hills landscape as the seventeenth participant was unavailable for an interview. This farm was excluded from the comparative analysis of farm management. Confounding variables were controlled by selecting farms that used uniform cultivation methods, such as manual tillage and rain-fed irrigation. Finally, interview data were cross-verified during each sampling session through direct field observations of farm infrastructure, such as the presence of compost pits versus discarded chemical packaging, and assessments of vegetation complexity.

### Landscape context analysis

At the landscape scale, using a 500 m radius buffer per site, highly intensified landscapes were defined as those where cropland comprised more than 70% of the total area; however, while cropland was dominant, the farms typically consisted of small, fragmented parcels surrounded by some trees or bushes at the edges of fields. Medium-intensity landscapes were characterised by a cropland cover of between 30% and 70%, while the least intensified landscapes contained less than 30% cropland cover. The remainder of the matrix across these landscape categories consisted of a mosaic of semi-natural habitats and alternative land-use types, specifically including montane forest, riverine forest, plantation forest, agroforestry systems, shrubland, thickets, grassland, sisal plantations, and built-up areas.

Landscape context was quantified through spatial analysis in ArcGIS pro version 3.5.3, utilising the high-resolution land-cover map [51]. This dataset is based on a comprehensive land-cover map of the Taita Hills region generated from 2020 imagery. The classification methodology integrated temporal metrics from Sentinel-1 and Sentinel-2 satellites with topoclimatic data from the NASA Shuttle Radar Topography Mission (SRTM). To ensure thematic reliability, these remote sensing layers were calibrated against an extensive reference database of field-mapped land cover and vegetation.

For each study plot, we generated a 500 m-radius buffer around the plot centre and calculated the percentage of the buffer area classified as Semi-Natural Habitat (SNH) and Edge Density in ArcGIS. While a multitude of landscape metrics are available in the ArcGIS environment, these two indicators were prioritised to characterise the fundamental dimensions of landscape composition and configuration while minimising collinearity and statistical redundancy.

Selecting a 500 m radius served as a strategic compromise between spatial independence and ecological relevance. Larger radii resulted in significant spatial overlap among study sites, obscuring discrete environmental differences, whereas smaller radii were deemed insufficient to capture the broad foraging ranges of mobile nocturnal pollinators. This scale is particularly relevant to hawkmoths, which are known to traverse large areas in search of resources.

## Data analysis

We investigated the effects of landscape composition, configuration, and farm management on *C. papaya* pollinator communities, with a focus on the abundance of diurnal and nocturnal visitors, hawkmoths, other moths, and bees. To account for the hierarchical structure of our data and the nature of ecological count data, we employed Generalised Linear Mixed Models (GLMMs) using the glmmTMB package in R [52].

Prior to model construction, we assessed multicollinearity among our landscape metrics at the 500m scale. Due to high redundancy between several metrics (|r| > 0.7), we selected semi-natural habitat (% SNH) as a measure of landscape composition and edge density as a measure of landscape configuration. To ensure the comparability of effect sizes and improve model convergence, we standardised all continuous explanatory variables, % SNH and edge density, by Z-transformation (scaling to a mean of zero and unit variance) [53].

Given evidence of significant overdispersion in our response variables, where the variance substantially exceeded the mean, we used Negative Binomial distribution (specifically the *nbinom2* parameterisation) for all models. This approach is more robust than standard Poisson models for capturing the variability common in pollinator visitation rates [54]. We included farm management [organic vs conventional], % SNH, edge density, male or female tree, and time of day (day vs night) as fixed effects. To account for repeated measures and our nested experimental design, we incorporated Plot ID and Tree ID as nested random effects, with Year included as an additional random factor to account for inter-annual variation.

We performed model validation using the *DHARMa* package [55]. We inspected simulated residuals to confirm the absence of overdispersion and zero-inflation and utilised the *performance* package to verify that Variance Inflation Factors (VIF) remained within acceptable limits (< 5). Model results were visualised by extracting predicted marginal effects with the *ggeffects* package [56], which enables interpretation on the original response scale while holding other covariates constant.

To evaluate how isolation from pollen sources influences reproductive output, we modelled fruit set (defined as a binary success/failure response) using a binomial generalised linear mixed-effects model (GLMM). Models were implemented using the lme4 package in R, with the distance to the nearest pollen-producing conspecific (male or hermaphrodite tree) designated as the primary explanatory variable.

To evaluate the influence of pollination services on papaya reproductive success, we analysed three key yield metrics: fruit set, fruit weight, and total production (calculated as the product of fruit set and average fruit weight per treatment). Statistical analyses were performed in R [57], primarily utilising the *lme4* package for mixed-effects modelling [58]. Plot ID was included as a random effect in all candidate models to account for environmental and spatial variation across study locations. This random structure was selected as it yielded a lower Akaike Information Criterion (AICc) [30,59] than alternative structures containing Tree ID or year. Rather than employing step-wise deletion on the fixed effects, we performed model averaging across this restricted candidate set to account for model selection uncertainty and ensure robust parameter estimates.

We modelled fruit set using Generalised Linear Mixed Models [GLMMs] with a Poisson distribution and log link function [55]. Model diagnostics were performed using the *performance* package [56]; Pearson’s Chi-squared tests confirmed the absence of overdispersion [dispersion ratio = 0.898, p = 0.641].

To determine the drivers of reproductive success, we compared models using the corrected AICc. A model accounting for the additive contributions of both diurnal and nocturnal pollination periods (representing unrestricted open pollination) provided a significantly better fit (ΔAICc > 2 compared to single-period models; full model AICc = 142.7). Estimated Marginal Means (EMMs) were calculated using the *emmeans* package to conduct post hoc pairwise comparisons [60].

Fruit weight and total production (yield) were analysed using Linear Mixed Models (LMMs) with restricted maximum likelihood (REML) estimation. Degrees of freedom and p-values for fixed effects were estimated using Satterthwaite’s method via the *lmerTest* package [61].

For fruit weight comparisons involving hand pollination, we applied the Holm adjustment to control the family-wise error rate [62]. For total production, we again utilised AICc to evaluate model complexity.

All yield data were visualised using the *ggplot2* [63] and *ggbeeswarm* [64] packages. We utilised boxplots to show the distribution of data (medians and interquartile ranges), overlaid with quasirandomly jittered points representing individual observations to ensure full data transparency. All plots used a consistent colour palette to distinguish between treatment groups: light blue for open pollination, yellow for day-only, grey for night-only, and white for closed controls.

To analyze the relationship between pollination activity and fruit set, we employed a Generalized Linear Model (GLM) with a negative binomial distribution to account for overdispersion in the count data in R. The model evaluated total fruits set, fruit weight, and yield as the response variables, with diurnal visitor, nocturnal visit, and farm management included as fixed explanatory variables, and Plot ID used as a random effect.

## Data availability

The primary data supporting the findings of this study, including pollinator visitation rates, fruit set measurements, and fruit quality metrics, are available from the corresponding author upon reasonable request. The landscape land-use data utilized in our analysis were derived from the high-resolution land-cover framework established by Abera et al. (2022). These data are openly available via the repository associated with the original mapping study (https://doi.org/10.3390/data7030036). This dataset integrates temporal metrics from satellite imagery with topoclimatic data, and the associated spatial analyses were performed using standard geographic information system software. These data are accessible through the repository associated with the original mapping study.

## Code availability

The R scripts and custom computational workflows utilised to replicate all statistical analyses, landscape modelling, and data visualisations reported in this study are available from the corresponding author upon reasonable request. The provided code includes full documentation to ensure the reproducibility of the generalised linear mixed models (GLMMs) and the niche partitioning analysis across the reported diel cycles. Future updates or additional versions of the analysis scripts may be obtained by contacting the authors directly.

## Acknowledgments

This research was supported by the Deutsche Forschungsgemeinschaft (DFG) (GZ: OT 634/2-1). We extend our deepest gratitude to the papaya farmers in the Taita Hills for their hospitality and for granting access to their orchards, without which this work would not have been possible. We thank Prof. Petri Pellikka for granting us access to the land cover dataset used in this study. We are particularly grateful to Mercy Kibii and Nelson Mgenyi for their invaluable assistance with field data collection and logistical coordination. We also thank the students from the University of Würzburg, Caroline, Liz, Luca, Marja, Sophia, and all the drivers and other staff at the field station in Taita Hills for their dedicated support in the field and for the research team. Institutional support and laboratory facilities were provided by the University of Embu, and we thank Mr. Morris Mutua from the National Museums of Kenya (NMK) for his expert assistance with insect taxonomy.

## Author Contribution Statement

Conceptualisation: MO, ISD, MP

Methodology: MO, ISD, MP, JZ, SMP

Investigation: MO, ISD, MP, SMP

Visualisation: MO, ISD, MP, JZ, SMP

Funding acquisition: MO, ISD, MP

Project administration: MO, ISD

Supervision: ISD

Writing – original draft: MO, ISD, MP

Writing – review and editing: MO, ISD, MP, JZ, SMP

## Supplementary materials

**Table S1:** Pairwise comparisons of papaya reproductive success across pollination treatments. Summary of post-hoc contrasts (Estimated Marginal Means) evaluating the differences between pollination modes for fruit set, fruit weight, and total production. For fruit set, values represent the ratio between treatments (calculated on the log scale), where a ratio >1 indicates higher success in the first group. For fruit weight (g) and production (kg), values represent the absolute mean difference (Estimate) between groups. All comparisons used the Kenward-Roger degrees-of-freedom method.

| Response Variable | Contrast | Ratio / Estimate | Std. Error | z/t ratio | P value |
| --- | --- | --- | --- | --- | --- |
| Fruit set (Ratio) | Open / Day pollination | 1.58 | 0.392 | 1.832 | 0.067 |
|  | Open / Night pollination | 2.05 | 0.554 | 2.654 | 0.008 |
|  | Open / Closed flowers | 20.5 | 14.7 | 4.207 | <0.001 |
|  | Day / Night pollination | 1.3 | 0.383 | 0.89 | 0.374 |
|  | Day / Closed flowers | 13 | 9.46 | 3.525 | <0.001 |
|  | Night / Closed flowers | 10 | 7.35 | 3.131 | 0.002 |
| Fruit weight (g) | Hand pollination – Open | 358 | 87.3 | 4.103 | <0.001 |
|  | Hand pollination – Closed | 1230 | 244 | 5.043 | <0.001 |
|  | Open – Day pollination | 202.5 | 108 | 1.871 | 0.064 |
|  | Open – Night pollination | 265.6 | 115 | 2.303 | 0.023 |
|  | Open – Closed flowers | 812.7 | 240 | 3.391 | 0.001 |
|  | Day – Night pollination | 63.1 | 124 | 0.507 | 0.613 |
|  | Day – Closed flowers | 610.3 | 247 | 2.475 | 0.0150 |
|  | Night - Closed flowers | 547.1 | 247 | 2.215 | 0.0291 |
| Yield (kg) | Open – Day pollination | 1.902 | 0.67 | 2.838 | 0.009 |
|  | Open – Night pollination | 2.579 | 0.67 | 3.848 | <0.001 |
|  | Open – Closed flowers | 4.1 | 0.67 | 6.118 | <0.001 |
|  | Day – Night pollination | 0.677 | 0.67 | 1.01 | 0.322 |
|  | Day – Closed flowers | 2.198 | 0.67 | 3.28 | 0.003 |
|  | Night – Closed flowers | 1.522 | 0.67 | 2.27 | 0.031 |

**Table S2:** Results of Negative Binomial Generalized Linear Models (GLM) assessing the effects of diurnal and nocturnal visitor abundance and farm management on total fruit set and average fruit weight.

| **Response Variable** | **Explanatory variable** | **Estimate** | **Std. Error** | **z/t value** | **P value** |
| --- | --- | --- | --- | --- | --- |
| Fruit set | Intercept | 3.2747 | 0.546 | 5.998 | <0.001 |
|  | Diurnal visitor abundance | -0.005 | 0.005 | -1.071 | 0.284 |
|  | Nocturnal visitor abundance | 0.001 | 0.004 | 0.28 | 0.779 |
|  | Farm management | -0.6668 | 0.652 | -1.022 | 0.307 |
| Fruit weight (g) | Intercept | 7.1372 | 0.528 | 13.515 | <0.001 |
|  | Diurnal visitor abundance | -0.0011 | 0.004 | -0.25 | 0.803 |
|  | Nocturnal visitor abundance | -0.0006 | 0.003 | -0.192 | 0.848 |
|  | Farm management | -0.0783 | 0.626 | -0.125 | 0.9 |
| Yield (kg) | Intercept | 3.2747 | 0.546 | 5.998 | <0.001 |
|  | Diurnal visitor abundance | -0.005 | 0.005 | -1.071 | 0.284 |
|  | Nocturnal visitor abundance | 0.001 | 0.004 | 0.28 | 0.779 |
|  | Farm management | -0.667 | 0.652 | -1.022 | 0.307 |

## References

1. Andres, C., & Bhullar, G. S. (2016). Sustainable intensification of tropical agro-ecosystems: Need and potentials. Frontiers in Environmental Science, 4, 5. 10.3389/fenvs.2016.00005

2. Power, A. G. (2010). Ecosystem services and agriculture: Tradeoffs and synergies. Philosophical Transactions of the Royal Society B: Biological Sciences, 365(1554), 2959–2971. 10.1098/rstb.2010.0143

3. Sena, V. G. L., de Moura, E. G., Macedo, V. R. A., Aguiar, C. F., Price, A. H., Mooney, S. J., & Calonego, J. C. (2020). Ecosystem services for intensification of agriculture, with emphasis on increased nitrogen ecological use efficiency. Ecosphere, 11(11), e03028. 10.1002/ecs2.3028

4. Tscharntke, T., Klein, A. M., Kruess, A., Steffan-Dewenter, I., & Thies, C. (2005). Landscape perspectives on agricultural intensification and biodiversity– ecosystem service management. Ecology Letters, 8(8), 857–874. 10.1111/j.1461-0248.2005.00782.x

5. Altendorf, S. (2019). Minor tropical fruits: Mainstreaming a niche market. Food Outlook, FAO.

6. Food and Agriculture Organization. (2023). Major tropical fruits market review 2022. FAO. https://openknowledge.fao.org/handle/20.500.14283/cc7108en

7. Kehlenbeck, K., Asaah, E., & Jamnadass, R. (2013). Diversity of indigenous fruit trees and their contribution to nutrition and livelihoods in Sub-Saharan Africa: Examples from Kenya and Cameroon. In Diversifying Food and Diets (pp. 257–269). Routledge.

8. Miller, D. C., Muñoz-Mora, J. C., & Christiaensen, L. (2020). Do trees on farms improve household well-being? Evidence from national panel data in Uganda. Frontiers in Forests and Global Change, 3, 101. 10.3389/ffgc.2020.00101

9. Kasina, J. M., Mburu, J., Kraemer, M., & Holm-Mueller, K. (2009). Economic valuation of animal pollination services: An assessment of various crops in Kakamega, Western Kenya. Journal of Economic Entomology, 102(3), 897–904. 10.1603/029.102.0307

10. Gemmill-Herren, B., Kwapong, P. K., Aidoo, K., Martins, D., Kinuthia, W., Gikungu, M., & Eardley, C. D. (2014). Priorities for Research and Development in the Management of Pollination Services for Agricultural Development in Africa. Journal of Pollination Ecology, 12, 40–51. 10.26786/1920-7603(2014)1

11. Stewart, R. I., Carvalheiro, L. G., Henson, I. W., Belcher, B. M., Bedano, J. C., Fedoroff, N. V., … & Firbank, L. G. (2014). The rest of the world is not the West: Trusting and managing ecosystem services in a changing tropical landscape. Philosophical Transactions of the Royal Society B: Biological Sciences, 369(1639), 20130232. 10.1098/rstb.2013.0232

12. Koul, B., Pudhuvai, B., Sharma, C., Kumar, A., Sharma, V., Yadav, D., & Jin, J.-O. (2022). Carica papaya L.: A tropical fruit with benefits beyond the tropics. Diversity, 14(8), 683. 10.3390/d14080683

13. Devoto, M., Bailey, S., Craze, P., & Memmott, J. (2011). Understanding and planning ecological restoration using network theory. Ecology Letters, 14(10), 1048–1061. 10.1111/j.1461-0248.2011.01673.x

14. Hahn, M., Scherer, G., & Linsenmair, K. E. (2021). Nocturnal pollinators: An overlooked component of biodiversity in tropical agricultural landscapes. Biotropica, 53(3), 742–754.

15. Macgregor, C. J., Pocock, M. J., Fox, R., & Evans, D. M. (2015). Pollination by nocturnal Lepidoptera, and the effects of light pollution: A review. Ecological Entomology, 40(3), 187–198. 10.1111/een.12174

16. Kremen, C., Iles, A., & Bacon, C. (2012). Diversified farming systems: An agroecological, systems-based alternative to modern industrial agriculture. Ecology and Society, 17(4), 44. 10.5751/ES-05103-170444

17. Lichtenberg, E. M., Kennedy, C. M., Kremen, C., Batáry, P., Berendse, F., Bommarco, R., … & Crowder, D. W. (2017). A global synthesis of the effects of diversified farming systems on arthropod diversity within fields and across agricultural landscapes. Global Change Biology, 23(11), 4946–4957. 10.1111/gcb.13714

18. Ávila Hernández, J. G., Cárdenas Aquino, M. d. R., Camas Reyes, A., & Martínez Antonio, A. (2023). Sex determination in papaya: Current status and perspectives. Plant Science, 335, 111814. 10.1016/j.plantsci.2023.111814

19. Parés Cascante, L., et al. (2024). Reproductive biology and pollination syndromes in the Caricaceae: A global review. Botanical Journal of the Linnean Society, 204(1), 45–62.

20. Dey, L., Mondal, S., & Mandal, S. (2016). Flower visitor diversity with reference to pollen dispersal and pollination of Carica papaya L. International Journal of Advanced Research in Biological Sciences, 3(2), 65–71.

21. Garibaldi, L. A., Steffan-Dewenter, I., Winfree, R., Aizen, M. A., Bommarco, R., Cunningham, S. A., … & Klein, A. M. (2013). Wild pollinators enhance fruit set of crops regardless of honey bee abundance. Science, 339(6127), 1608–1611. 10.1126/science.1230200

22. Badillo-Montaño, R., Aguirre, A., & Munguía-Rosas, M. A. (2018). Pollinator-mediated interactions between cultivated papaya and co-flowering plant species. Ecology and Evolution, 9(1), 587–597. 10.1002/ece3.4781

23. Renner, S. S., & Feil, J. P. (1993). Pollinators of tropical dioecious angiosperms. American Journal of Botany, 80(9), 1100–1107. 10.1002/j.1537-2197.1993.tb15337.x

24. Martins, D. J., & Johnson, S. D. (2009). Distance and quality of natural habitat influence hawkmoth pollination of cultivated papaya. International Journal of Tropical Insect Science, 29(3), 114–123. 10.1017/S1742758409990208

25. Ricketts, T. H., Regetz, J., Steffan-Dewenter, I., Cunningham, S. A., Kremen, C., Bogdanski, A., … & Winfree, R. (2008). Landscape effects on crop pollination services: a global synthesis. Ecology Letters, 11(5), 499–515. 10.1111/j.1461-0248.2008.01157.x

26. Kremen, C., Williams, N. M., Aizen, M. A., Gemmill-Herren, B., LeBuhn, G., Minckley, R., … & Winfree, R. (2007). Pollination and other ecosystem services produced by mobile organisms: a conceptual framework for the effects of land-use change. Ecology Letters, 10(4), 299–314. 10.1111/j.1461-0248.2007.01018.x

27. Wright, H. J., Eardley, C. D., Kühn, I., & Penev, L. (2015). Priorities for research and development in the management of pollination services for agriculture in Africa. Journal of Pollination Ecology, 15(1), 1–11.

28. Aizen, M. A., Garibaldi, L. A., Cunningham, S. A., & Klein, A. M. (2008). Long-term global trends in crop yield and production reveal no current pollination shortage but increasing ecology of reliance. Current Biology, 18(20), 1572–1575. 10.1016/j.cub.2008.08.066

29. Willmer, P. (2011). Pollination and floral ecology. Princeton University Press.

30. Khalifa, S. A. M., Elshafiey, E. H., Shetaia, A. A., El Wahed, A. A. A., Algethami, A. F., Musharraf, S. G., AlAjmi, M. F., Zhao, C., Masry, S. H. D., Abdel Daim, M. M., Halabi, M. F., Kai, G., Al Naggar, Y., Bishr, M., Diab, M. A. M., & El Seedi, H. R. (2021). Overview of bee pollination and its economic value for crop production. Insects, 12(8), 688. 10.3390/insects12080688

31. Kendall, L., & Nicholson, C. C. (2025). Pollination Across the Diel Cycle: A Global Meta-Analysis. Ecology Letters, 28, e70036. 10.1111/ele.70036

32. Eardley, C., & Urban, R. (2010). Catalogue of Afrotropical bees (Hymenoptera: Apoidea: Apiformes). Zootaxa, 2455(1), 1–548. 10.11646/zootaxa.2455.1.1

33. Dafni, A. (1984). Mimicry and deception in pollination. Annual Review of Ecology and Systematics, 15, 259–278. 10.1146/annurev.es.15.110184.001355

34. Vereecken NJ, Schiestl FP. The evolution of imperfect floral mimicry. Proc Natl Acad Sci U S A. 2008 May 27;105(21):7484–8. 10.1073/pnas.0800194105

35. Raguso, R.A., & Pellmyr, O. (1998). Dynamic headspace analysis of floral volatiles: A comparison of methods. Oikos, 81, 238–254. 10.2307/3547045.

36. Parthasarathy K, Willis MA. Spatial odor discrimination in the hawkmoth, Manduca sexta (L.). Biol Open. 2021 Mar 26;10(3):bio058649. doi: 10.1242/bio.058649.

37. Jürgens A, Dötterl S, Meve U. The chemical nature of fetid floral odours in stapeliads (Apocynaceae-Asclepiadoideae-Ceropegieae). New Phytol. 2006;172(3):452–68. doi: 10.1111/j.1469-8137.2006.01845.x.

38. Boreux, V., Kushalappa, C. G., Vaast, P., & Ghazoul, J. (2013). Interactive effects of landscape context and management on pollination services in a coffee agroforestry system. Landscape Ecology, 28(8), 1529–1543. 10.1007/s10980-013-9906-z

39. Stöckl, A. L., & Kelber, A. (2019). Fuelling on the wing: Sensory ecology of hawkmoth foraging. Journal of Comparative Physiology A, 205(3), 399–413. 10.1007/s00359-019-01328-2

40. Klein, A.-M., Vaissière, B. E., Cane, J. H., Steffan Dewenter, I., Cunningham, S. A., Kremen, C., & Tscharntke, T. (2007). Importance of pollinators in changing landscapes for world crops. Proceedings of the Royal Society B: Biological Sciences, 274(1608), 303–313. 10.1098/rspb.2006.3721

41. Andrieu, E., Vialatte, A., Sirami, C., & Balent, G. (2007). Local and landscape effects on bees in Mediterranean fragmented landscapes. Agriculture, Ecosystems & Environment, 120(2–4), 213–221. 10.1016/j.agee.2006.09.004

42. Fahrig, L., Baudry, J., Brotons, L., Burel, F. G., Crist, T. O., Fuller, R. J., … & Martin, J. L. (2011). Functional heterogeneity in agricultural landscapes for biodiversity conservation. Ecology Letters, 14(2), 101–112. 10.1111/j.1461-0248.2010.01559.x

43. Wiens, J. A. (1989). Spatial scaling in ecology. Functional Ecology, 3(4), 385–397. 10.2307/2389612

44. Hole, D. G., Perkins, A. J., Wilson, J. D., Alexander, I. H., Grice, P. V., & Evans, A. D. (2005). Does organic farming benefit biodiversity? Biological Conservation, 122(1), 113–130. 10.1016/j.biocon.2004.07.018

45. Rundlöf, M., Andersson, G. K., Bommarco, R., Fries, I., Hederström, V., Herbertsson, L., Jonsson, O., Klatt, B. K., Pedersen, T. R., Yourstone, J., & Smith, H. G. (2015). Seed coating with a neonicotinoid insecticide negatively affects wild bees. Nature, 521(7550), 77–80. 10.1038/nature14420

46. Otieno, M., Peters, M. K., & Steffan Dewenter, I. (2025). Safeguarding nocturnal pollinators for food security in sub Saharan Africa. Trends in Ecology & Evolution, 40(10), 931–934. 10.1016/j.tree.2025.08.002

47. Johnson, S. D., Moré, M., Amorim, F. W., Haber, W. A., Frankie, G. W., Stanley, D. A., Cocucci, A. A., & Raguso, R. A. (2017). The long and the short of it: A global analysis of hawkmoth pollination niches and interaction networks. Functional Ecology, 31, 101–115. 10.1111/1365-2435.12753

48. Rader, R., Bartomeus, I., Garibaldi, L. A., Garratt, M. P., Howlett, B. G., Winfree, R., Cunningham, S. A., Mayfield, M. M., Arthur, A. D., Andersson, G. K., Bommarco, R., Brittain, C., Carvalheiro, L. G., Chacoff, N. P., Entling, M. H., Foully, B., Freitas, B. M., Gemmill Herren, B., Ghazoul, J., … Woyciechowski, M. (2016). Non bee insects are important contributors to global crop pollination. Proceedings of the National Academy of Sciences of the United States of America, 113(1), 146–151. 10.1073/pnas.1517092112

49. Dzekashu, F. F., Yusuf, A. A., Pirk, C. W. W., Steffan-Dewenter, I., Lattorff, H. M. G., & Peters, M. K. (2022). Floral Turnover and Climate Drive Seasonal Bee Diversity Along a Tropical Elevation Gradient. Ecosphere, 13(3), e3964. 10.1002/ecs2.3964

50. Martins, D. J., & Johnson, S. (2013). Interactions between hawkmoths and flowering plants in East Africa: Polyphagy and evolutionary specialization in an ecological context. Biological Journal of the Linnean Society, 110(1), 119–140. 10.1111/bij.12107

51. Abera, T. A., Vuorinne, I., Munyao, M., Pellikka, P. K. E., & Heiskanen, J. (2022). Land cover map for multifunctional landscapes of Taita Taveta County, Kenya, based on Sentinel-1 radar, Sentinel-2 optical, and topoclimatic data. Data, 7(3), 36. 10.3390/data7030036

52. Brooks, M. E., Kristensen, K., van Benthem, K. J., Magnusson, A., Berg, C. W., Nielsen, A., Skaug, H. J., Maechler, M., & Bolker, B. M. (2017). glmmTMB balances speed and flexibility among packages for zero inflated generalized linear mixed modeling. The R Journal, 9(2), 378–400. 10.32614/RJ-2017-066

53. Schielzeth, H. (2010). Simple means to improve the interpretability of regression coefficients. Methods in Ecology and Evolution, 1(2), 103–113. 10.1111/j.2041-210X.2010.00012.x

54. Bolker, B. M., Brooks, M. E., Clark, C. J., Geange, S. W., Poulsen, J. R., Stevens, M. H. H., & White, J. S. S. (2009). Generalized linear mixed models: A practical guide for ecology and evolution. Trends in Ecology & Evolution, 24(3), 127–135. 10.1016/j.tree.2008.10.008

55. Hartig, F. (2022). DHARMa: Residual diagnostics for hierarchical (multi level / mixed) regression models (R package version 0.4.6).

56. Lüdecke, D. (2018). ggeffects: Tidy data frames of marginal effects from regression models. Journal of Open Source Software, 3(26), 772. 10.21105/joss.00772

57. R Core Team. (2026). R: A language and environment for statistical computing. R Foundation for Statistical Computing.

58. Bates, D., Mächler, M., Bolker, B., & Walker, S. (2015). Fitting linear mixed effects models using lme4. Journal of Statistical Software, 67(1), 1–48. 10.18637/jss.v067.i01

59. Ne’eman, G., Jürgens, A., Newstrom-Lloyd, L., Potts, S. G., & Dafni, A. (2010). A framework for comparing pollinator performance on multiple spatial scales. Biological Reviews, 85(2), 377–389. 10.1111/j.1469-185X.2009.00105.x

60. Lenth, R. V. (2023). emmeans: Estimated marginal means, aka least squares means (R package version 1.8.9).

61. Kuznetsova, A., Brockhoff, P. B., & Christensen, R. H. B. (2017). lmerTest package: Tests in linear mixed effects models. Journal of Statistical Software, 82(13), 1–26. 10.18637/jss.v082.i13

62. Holm, S. (1979). A simple sequentially rejective multiple test procedure. Scandinavian Journal of Statistics, 6(2), 65–70.

63. Wickham, H. (2016). ggplot2: Elegant graphics for data analysis. Springer. 10.1007/978-3-319-24277-4

64. Clarke, E., Sherrill Mix, S., & Dawson, C. (2023). ggbeeswarm: Categorical scatter plots with ggplot2 (R package version 0.7.2).

